# Tissue dynamics of the forebrain neural plate

**DOI:** 10.1101/016303

**Authors:** Stephen Young, Joel N. Jennings, Guy B. Blanchard, Alexandre J. Kabla, Richard J. Adams

## Abstract

The forebrain has the most complex shape and structure of the vertebrate brain regions and the mechanisms of its formation remain obscure. Convergence and extension movements are characteristic of the posterior (spinal cord and hindbrain) neural plate (pNP) while tissue deformations and underlying cellular dynamics during the early shaping of the forebrain neural plate (fNP) are undefined. Here, we apply live imaging, automated cell tracking and computational analysis to quantitatively map cell behaviour in the zebrafish fNP. We demonstrate a novel mechanism in which planar cell rearrangements, with a passive signature, are orthogonal to those in the pNP, and cell divisions lacking planar-polarity facilitate thickening from two to three layers. We develop a mechanical model of the fNP in which polarised cell behaviour arises from interactions with dissimilar bordering tissues rather than from intrinsically polarised cells. The model unifies *in vivo* observations and provides a mechanistic understanding of fNP morphogenesis.

## Introduction

The developmental processes shaping the vertebrate brain are a long-standing source of intrigue. Early in embryogenesis, neural tissues undergo complex sequential deformations. We are increasingly able to visualise these tissue dynamics in living embryos (Hirose et al., 2004; England et al., 2006; Rembold et al., 2006; Keller et al., 2008; Keller, 2013), and even to generate synthetic neural tissues *in vitro* that recapitulate a subset of them (Eiraku et al., 2011). Yet our understanding of their essential geometry and underlying mechanisms *in vivo* remains very incomplete. The spectrum of debilitating illnesses arising from failure of brain morphogenesis in humans provides a strong incentive for further conceptual progress.

During early vertebrate development, neurulation converts a sheet of neural progenitor cells, the neural plate, into the neural tube, a precursor structure possessing the major topographic features of the brain and spinal cord. The neural tube exhibits marked variations in shape along the head-tail axis because neurulation is not a single homogeneous process. Fate mapping of teleost, avian and mammalian embryos reveal evolutionarily conserved large-scale tissue rearrangements that are unique to forebrain neurulation; in particular, the fission of an initially continuous eye field to form bilateral retinae, and the anterior translocation of midline hypothalamic tissue beneath the plane of the retinae (Woo and Fraser, 1995; Varga et al., 1999; Lawson et al., 1999; Dale et al., 1999; Inoue et al., 2000; Cobos et al., 2001; Hirose et al., 2004; England et al., 2006). England et al. (2006) defined morphogenetic phases of forebrain neurulation by tracking the movement of cell populations within forebrain subdomains in the zebrafish embryo. Even during the earliest ‘contraction phase’, forebrain tissue deformations differed significantly from concurrent deformations in other domains of the neural plate.

Spinal cord neurulation lacks the large-scale tissue rearrangements of the forebrain and has received greater attention to date. Medio-lateral (ML) tissue convergence and anteroposterior (AP) tissue extension reflect the summation of multiple oriented cellular behaviours within the spinal cord neural plate that include planar cell intercalation (Jacobson and Gordon, 1976; Schoenwolf and Alvarez, 1992; Kimmel et al., 1994; Keller et al., 2000; Concha and Adams, 1998; Blanchard et al., 2009; Nishimura et al., 2012; Shindo and Wallingford, 2014), radial cell intercalation (Warga and Kimmel, 1990; Wilson et al., 1995), cell division (Schoenwolf and Alvarez, 1989; Kimmel et al., 1994; Sausedo et al., 1997; Concha and Adams, 1998; Gong et al., 2004; Quesada-Hernàndez et al., 2010), and cell shape changes (Papan and Campos-Ortega, 1994; Concha and Adams, 1998; Jessen et al., 2002; Blanchard et al., 2009). Planar cell polarity (non-canonical Wnt) signaling directly and indirectly controls the orientation of these inter-related cell behaviours (Wallingford et al., 2002; Roszco et al., 2009).

Initial observations of amphibian fNPs revealed no obvious planar cell intercalation (Burnside et al., 1968; Jacobson and Gordon, 1976; Elul et al., 1997), and cell divisions of surface layer cells in the zebrafish fNP are not oriented within the plane (Concha and Adams, 1998). These findings suggest either that fNP cells are not intrinsically planar-polarised, or that planar cell polarity is not deployed to orient mitosis and motile behaviour as seen in the pNP. fNP cells in the newt progressively lose cross-sectional area and grow taller *in vivo*, and surgical isolation of neural plates demonstrates that cells contract actively (Jacobson and Gordon, 1976). However, the apparent complexity of cell displacement fields in the early fNP suggests that cell contraction alone is insufficient to explain fNP morphogenesis. To address this question, we collected high-resolution time-lapse recordings of the zebrafish fNP and applied quantitative analyses to dissect the morphogenetic process at multiple scales. We detected a set of novel tissue behaviours in the fNP relative to the spinal cord and hindbrain neural plate. In addition, we have formulated a conceptual model to unify our observations and implemented this model as a physical simulation that provides insight into underlying tissue mechanics.

## Results

### Quantitative analysis of fNP tissue dynamics

To elucidate the mechanisms that drive morphogenesis of the forebrain, we performed quantitative analyses of tissue deformation and cellular behaviour in the zebrafish fNP (morphogenetic data are collated in Supplementary Table 1 and Supplementary Table 2). We focus on the first ‘contraction phase’ (England et al., 2006) of fNP neurulation, that takes place over a one-hour period beginning at ~8 hours post fertilisation (hpf) (Figure 1 and Supplementary Movie 1). We tracked the positions and shapes of fNP cells with labelled plasma membranes and nuclei (see Methods). At 8 hpf, the fNP is bilayered (Fig. 2e), with a distinct surface layer of discoidal cells and deep layer of cuboidal cells (Supplementary Figs. 1 and 2). Surface and deep layer deformations are comparable in most respects (Figs. 2g and 3d–h, and Supplementary Fig. 3).

**Figure 1.**
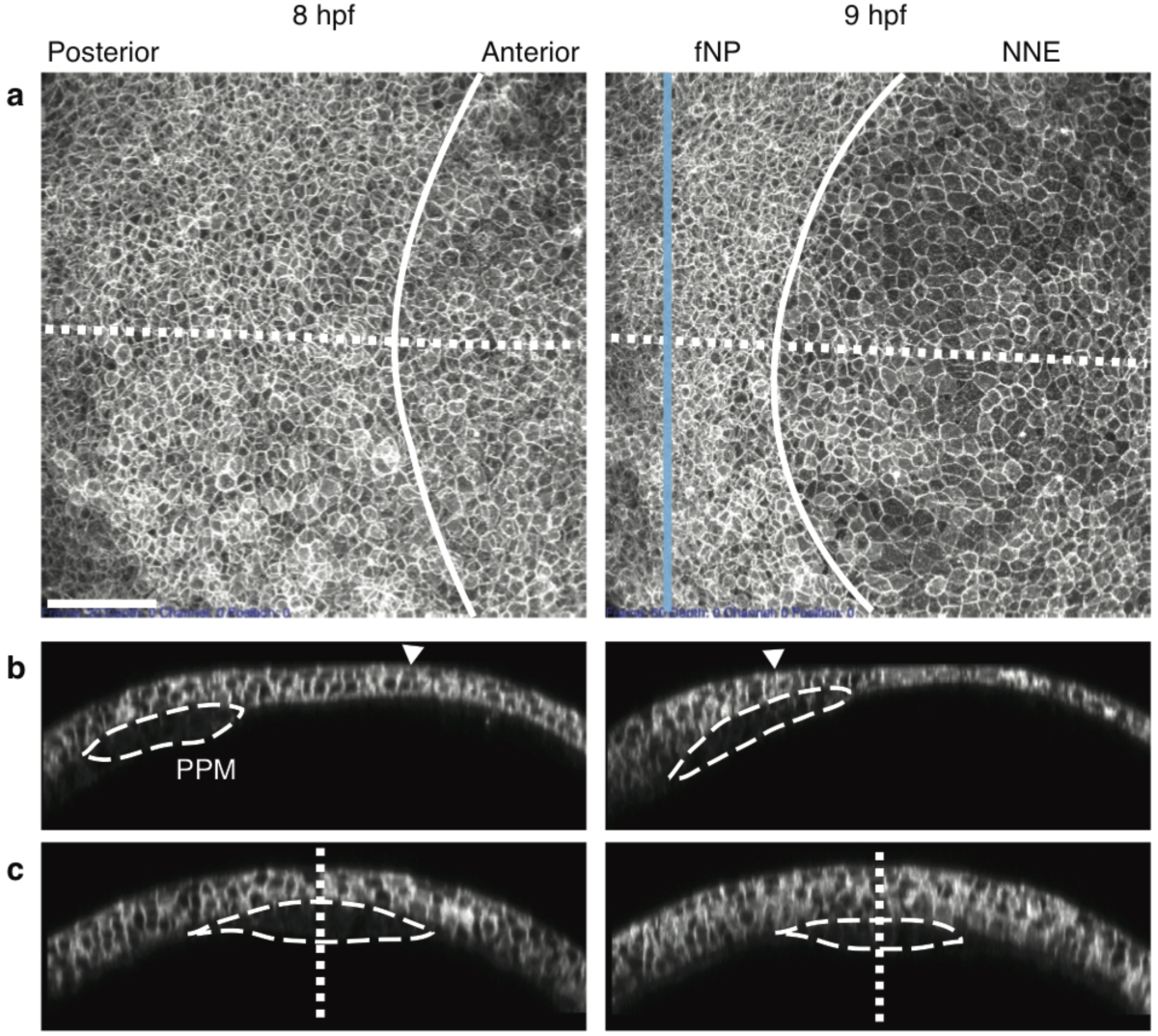
The contraction phase of the fNP (see Supplementary Movie 1). (**a**) Dorsal maximum intensity projection from a time-lapse confocal series. (**b**) Sagittal section through embryo midline. (**c)** Transverse sections indicated in (**a**) through the fNP. Solid line, neural/non-neural boundary. Dotted line, embryo midline. Dashed line, projection of PPM. Scale, 100 µm.

**Figure 2.**
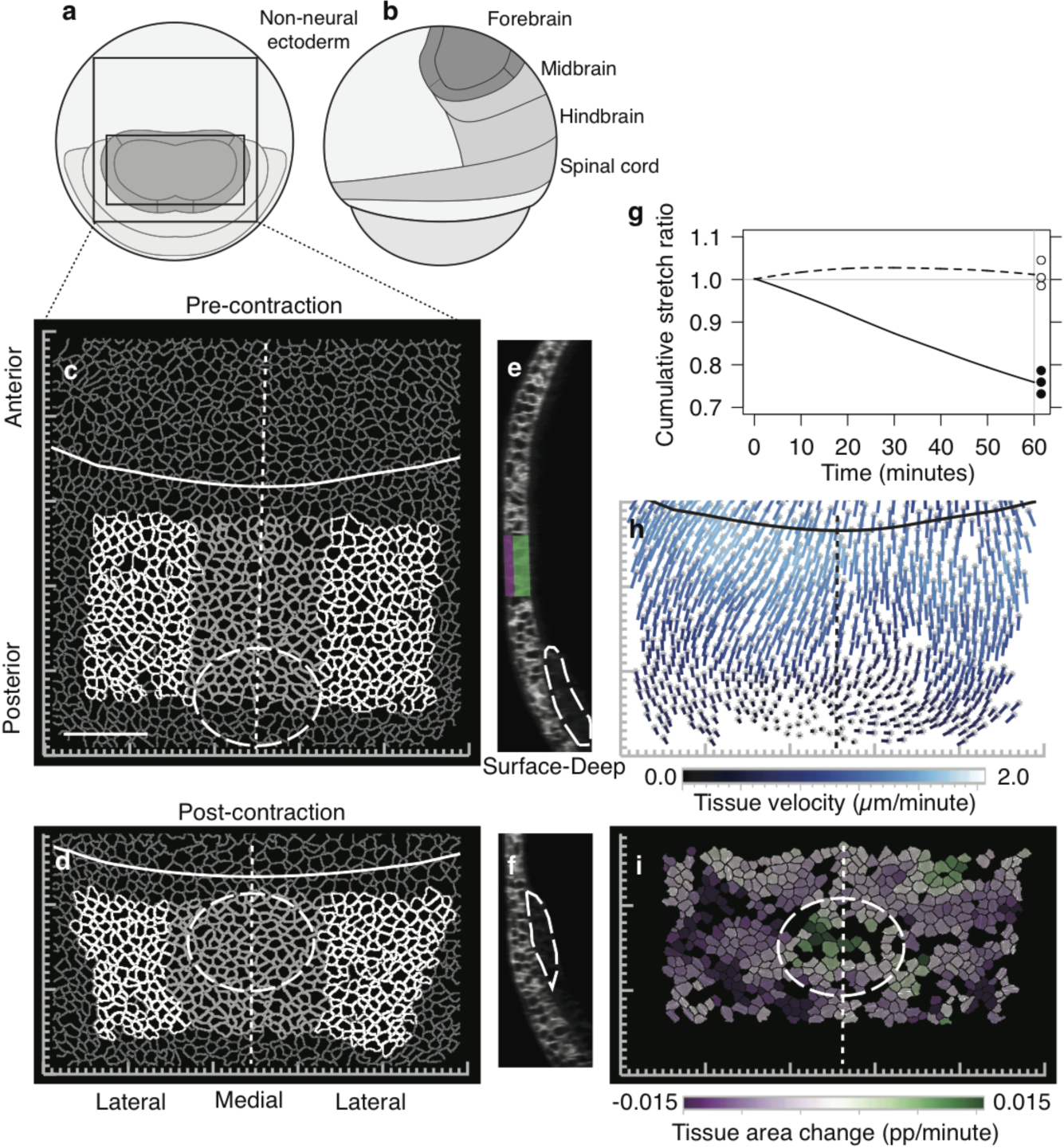
The early fNP undergoes anisotropic contraction. (**a**) Animal pole and (**b**) lateral view of the zebrafish ectoderm fate map (8 hpf) (Woo and Fraser, 1995; Schier and Talbot, 2005; England et al., 2006). (**c,d**) Tracked cell outlines in the ectoderm deep cell layer pre-(**c**) and post-contraction (**d**) (lateral fNP, white; medial fNP, light gray). (**e,f**) Confocal section through embryonic midline (surface layer, magenta; deep layer, green). (**g**) Tissue cumulative stretch ratio throughout the contraction phase for the entire fNP (solid line, AP; dashed line, ML). Filled circles, AP endpoints; open circles, ML endpoints, for 3 replicate embryos. (**h**) Local tissue velocity field (mid-contraction). Gray symbols mark initial location of tissue domains (**i**) Local planar tissue area change (mid-contraction). Solid line, neural/non-neural boundary. Dotted line, midline. Dashed line, projection of PPM. Scale, 100µm.

**Figure 3.**
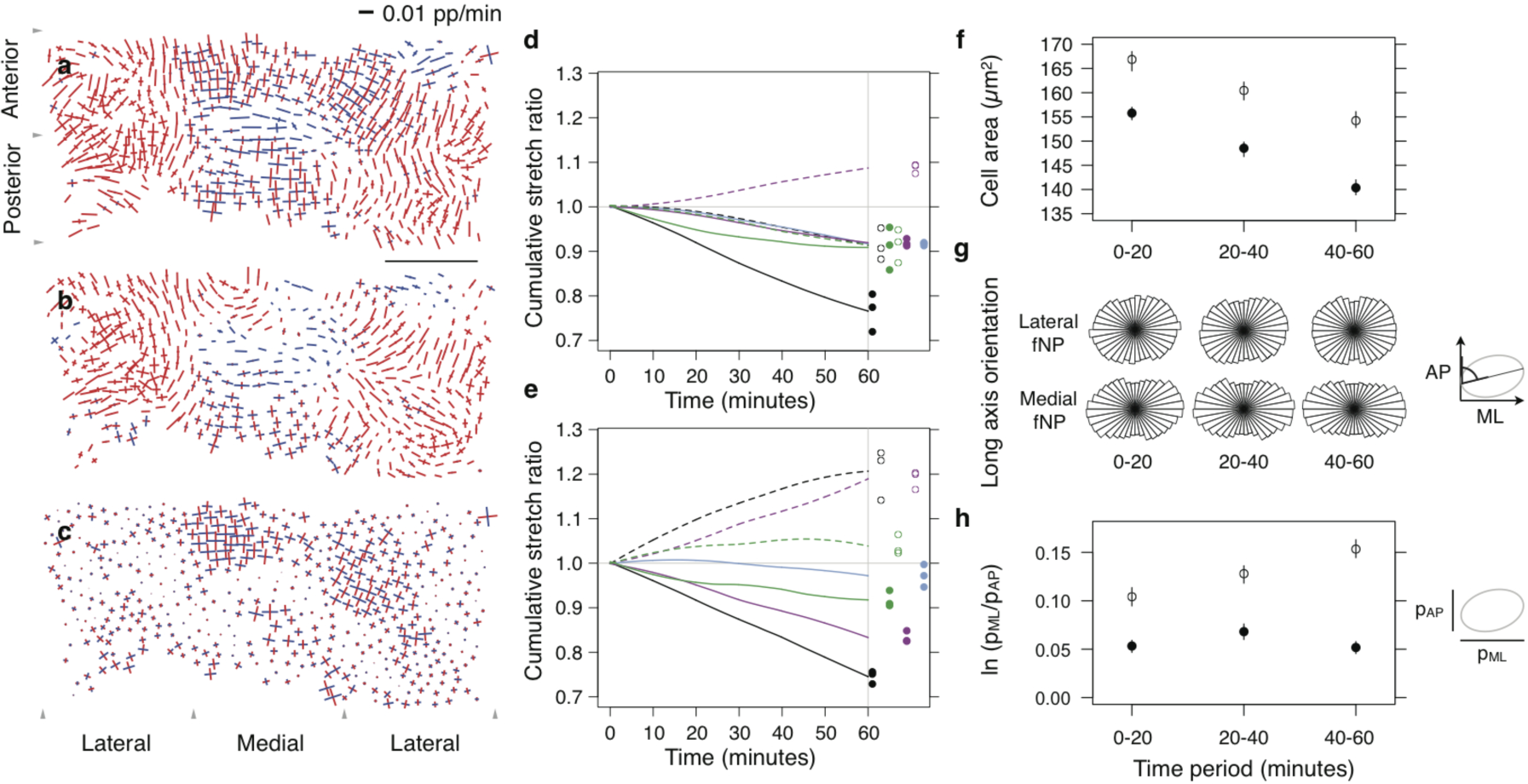
Regional cell deformation and intercalation processes reshape the fNP. (**a**-**c**) Tissue (**a**), cell shape (**b**) and cell intercalation (**c**) strain rate maps mid-contraction. Line segments show local orthogonal principal strain rates where length is equal to strain rate amplitude (blue, expansion; red, contraction) (Blanchard et al., 2009, see Methods). (**d,e**) Cumulative stretch ratios integrated across AP (solid) and ML (dotted) orientations for total tissue (black), cell shape (green), cell intercalation (magenta) and radial rearrangement (blue) in lateral (**d**) and medial (**e**) fNP regions. Filled circles, AP endpoints; open circles, ML endpoints, for 3 replicate embryos. (**f**) Cell area. (**g**) Cell long axis orientation. (**h**) Projected cell aspect ratio. (**f,h**) median, bars 95% confidence interval. Filled circles, lateral fNP; open circles, medial fNP. Scale, 100 µm.

### Planar contraction of the fNP is anisotropic and patterned

The planar area of the fNP decreases overall by ~25% during the contraction phase. This change is anisotropic; anteroposterior (AP) length is reduced by ~25% while the mediolateral (ML) width remains essentially constant (Fig. 1c, d and g). However, fNP subdomains exhibit distinct deformations, as seen by non-uniform cell trajectories (Fig. 2h). In particular, qualitatively-distinct lateral and medial ML deformations were observed; lateral regions contract ~8%, while the medial region undergoes a net extension of ~20% (Fig. 3a–e).

The medial fNP region exhibits a territory of planar area expansion that translates posterior to anterior through the fNP (Fig. 2i and Supplementary Movie 2). This feature correlates with the position of the underlying prechordal plate mesoderm (PPM), which moves anteriorly through directed migration, in direct contact with the medial fNP (Ulrich et al., 2003) (Fig. 2, d and f). Notably, the behaviour of medial fNP subregions prior to and after contact with the PPM is comparable to that of the lateral fNP (Supplementary Fig. 4 and Supplementary Movie 2). This suggests that the PPM induces a transient deviation from an otherwise global morphogenetic behaviour expressed across the fNP.

### fNP cells undergo progressive columnarisation

To investigate how local cell behaviours lead to anisotropic and heterogeneous tissue deformations, we decomposed tissue deformation into component contributions due to cell shape changes, cell intercalation (Blanchard et al., 2009) and cell division (Fig. 3a–e and Supplementary Tables 1 and 2) (see Methods).

Lateral fNP cells contract isotropically by ~8% along AP and ML axes (Fig. 3d), with areas decreasing by ~14%. Initially, cells have a modest 5% ML elongation (Fig. 3h) but ML contraction lags AP contraction so that after 30 minutes cells transiently reach 7% elongation (Fig. 3, d and h), with their long axes biased to the ML axis (Fig. 3g and Supplementary Table 3), before returning to their original 5% elongation (Fig. 3h).

Medial fNP cells on average decrease 11% in area over the contraction phase (Fig. 3f). They have more elongated shapes (Fig. 3h) with strong ML orientation (Fig. 3g and Supplementary Table 3). Cell deformations are modulated within the medial fNP overlaying the PPM (Supplementary Fig. 4). Cells within the posterior medial fNP initially expand but resume contraction once the PPM moves beyond them. Cells within the anterior medial fNP initiate planar contraction and then expand as the PPM advances underneath.

We propose that planar contraction is the intrinsic default cell behaviour expressed throughout the fNP. PPM migration beneath the medial fNP induces a local reversal of cellular deformations resulting in cell flattening and transient planar tissue expansion. This patterned morphogenesis does not reflect known variations in gene expression within the neural plate but more likely reflects the transient physical interaction between the two overlying tissues.

### fNP thickening and lamination by mitosis-associated radial cell rearrangement

Tissue area loss within the plane is balanced by radial thickening of the neural plate. In the lateral fNP, thickening is continuous throughout the contraction phase (Fig. 4a), whereas medial fNP regions transiently overlying the PPM show stalled thickening (Fig. 4b) or thinning (Fig. 4c). Approximately 50% of planar area reduction in the lateral fNP is attributable to cell shape changes; radial cell rearrangement accounts for the remainder (Fig. 3, d and e, and Fig. 5, a and b).

**Figure 4.**
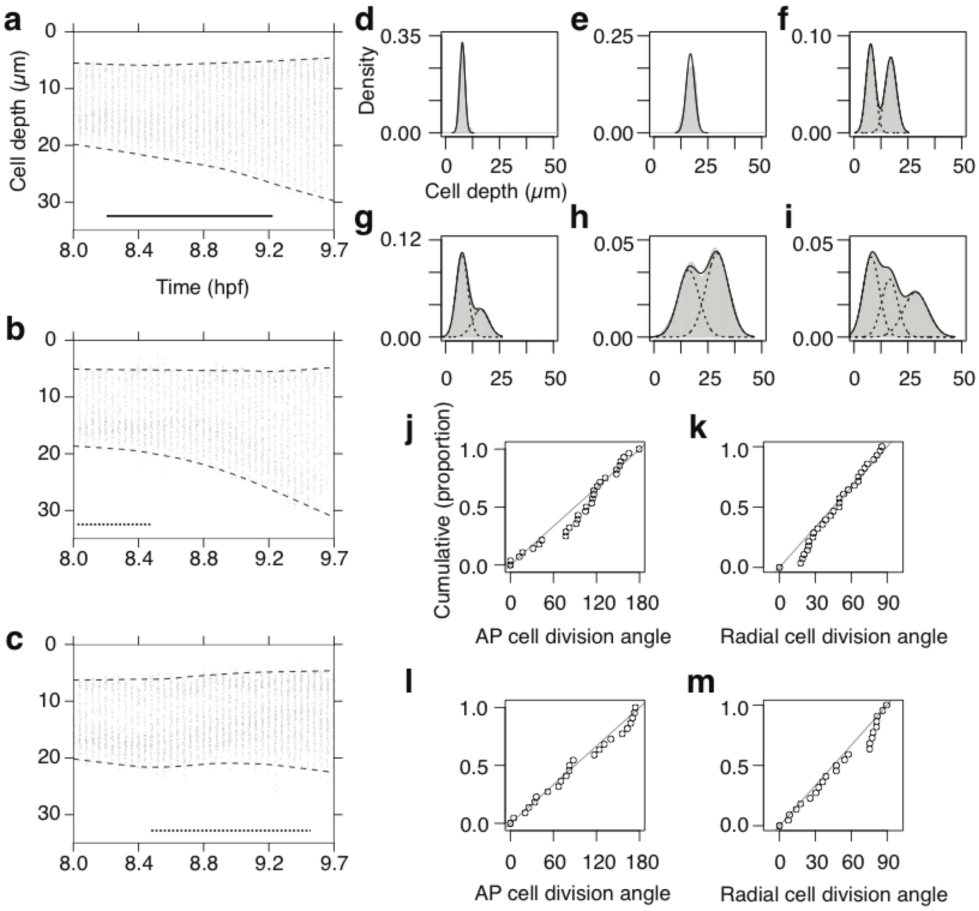
Radial cell rearrangement thickens the fNP and alters its layer structure. (**a-c**) Radial cell depth distribution time course for lateral (**a**), posterior medial (**b**) and anterior medial fNP (**c**) tissue. Dashed lines, 1st and 99th percentile; solid line, 60 minute contraction period in Figs. 1, 2 and 4. Dotted lines show time periods during which the PPM underlies the fNP region (**d** to **i**) Radial cell depth distributions at 8.0 hpf (pre-contraction) (**d**-**f**) and 9.7 hpf (post-contraction) (**g**-**i**) for cell populations originating in the lateral surface layer (**d,g**), lateral deep layer (**e,h**) or both layers (**f,i**), and their descendants. Solid gray area, kernel density estimate; solid line, sum of Gaussian fit; dashed line, component Gaussian curves. (**j**-**m**) Orientation of deep layer cell divisions in lateral (**j,k**) and medial (**l,m**) fNP relative to the AP (**j,l**) or radial axis (**k,m**) of the embryo. Distributions are not distinct from random at p>0.05 using Watson’s test for circular uniformity. Diagonal line, expected cumulative distribution for randomly oriented divisions.

**Figure 5.**
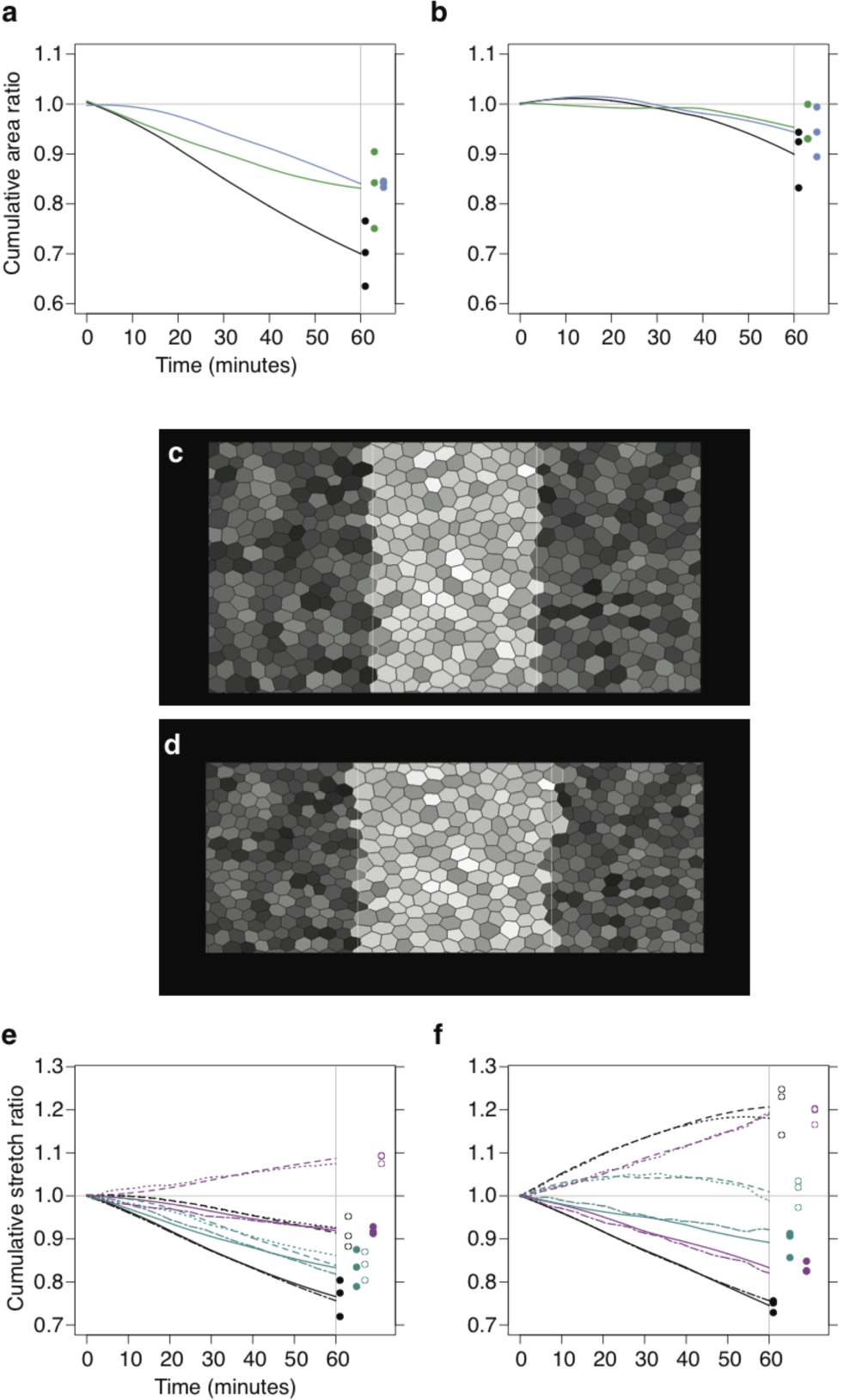
Isotropic active behaviour and passive responses to mechanical constraints can account for *in vivo* tissue dynamics. (**a,b)** Tissue area time course for lateral (**a**) and medial (**b**) fNP (black), and contribution from cell shape (green) and radial cell rearrangement (blue). Filled circles, endpoints for 3 replicate embryos. (**c,d**) Simulation snapshot at 0 (**c**) and 60 (**d**) minutes. Dark grays randomly colour lateral cells; light grays randomly colour medial cells (cells retain assigned colour during simulation). (**e,f**) Cumulative stretch ratios integrated across AP (experimental, solid; simulated, twodash) and ML (experimental, dashed; simulated, dotted) orientations for tissue (black), combined cell shape and radial rearrangement (dark cyan), and cell intercalation (magenta) in lateral (**e**) and medial (**f**) fNP regions. Filled circles, AP endpoints; open circles, ML endpoints, for 3 replicate embryos.

Radial cell rearrangement alters the layer structure of the fNP (Fig. 4, d–i, and Supplementary Table 1). The initial bilayer organisation (Figs. 2e and 4f) evolves as cells are displaced from both surface and deep layers (Fig. 4d, e, g and h) to create an intermediate population of cells separated from the tissue surfaces (Fig 4i). This process appears unique to fNP. In the spinal cord and hindbrain neural plate, radial cell intercalation occurs in the opposite direction, collapsing cell layers and effecting planar tissue expansion and radial thinning (Keller et al., 1992; Keller et al., 2000; Hong and Brewster, 2006).

### fNP cell divisions are not planar-polarised

An asynchronous round of cell divisions occurs within the fNP during the contraction phase (Kimmel et al., 1994; Concha and Adams, 1998). Most cells forming the intermediate population are daughter cells displaced during or soon after cell division. However, despite a consistent orientation of radial cell rearrangement, cell division orientation itself is random in the deep layer of the fNP; divisions are neither biased relative to the tissue plane nor oriented within the plane (Fig. 4j–m, and Supplementary Table 3). Surface layer divisions in the lateral and medial fNP show a clear bias toward planar orientations, but no consistent orientation within the plane (Concha and Adams, 1998) (Supplementary Fig. 5 and Supplementary Table 3). This contrasts markedly with the early zebrafish spinal cord neural plate, where cell divisions are strongly biased to planar and AP orientations (Concha and Adams, 1998; Gong et al., 2004). Interestingly, cell divisions in spinal cord tissue in which planar cell polarity signaling has been disrupted are neither biased relative to the tissue plane nor oriented within the plane (Gong et al., 2004), as shown here for the deep layer of the fNP.

### fNP exhibits reversed planar intercalation relative to trunk neural tissue

Radial dispersion of daughter cells thus represents a second intrinsic mechanism effecting planar contraction of the tissue, in addition to the ongoing planar cell shape contraction. However, neither of these morphogenetic processes show the planar anisotropy required to account for the observed changes in the aspect ratio of the tissue.

Tissue convergence and extension associated with oriented cell intercalation has been observed in the spinal cord and hindbrain neural plate of all vertebrates examined (Burnside and Jacobson, 1968; Jacobson and Gordon, 1976; Schoenwolf and Alvarez, 1989; Keller et al., 1992; Kimmel et al., 1994; Elul et al., 1997; Concha and Adams, 1998; Hirose et al., 2004; Hong and Brewster, 2006; Ybot Gonzalez et al., 2007; Blanchard et al., 2009). In contrast, cell dispersion and cell tracking assays indicated that cell rearrangement is limited or absent in the fNP (Burnside and Jacobson, 1968; Jacobson and Gordon, 1976; Elul et al., 1997; Hirose et al., 2004). We detected a global process of planar cell intercalation that widens the ML tissue axis while contracting the AP tissue axis (Fig. 3c–e). This is the inverse of rearrangements taking place concurrently in the spinal cord and hindbrain neural plate.

Cell intercalation in the spinal cord and hindbrain is thought to arise through an active process of oriented protrusion and contraction of adhesive subcellular extensions (Keller et al., 1992; Concha and Adams, 1998; Keller et al., 2000; Wallingford et al., 2002; Hong and Brewster, 2006; Brodland, 2006), or by the contraction of myosin-rich ML-oriented junctions (Nishimura et al., 2012; Shindo and Wallingford, 2014). We infer that planar intercalation in the zebrafish fNP is instead a passive morphogenetic process. When active cell intercalation drives tissue convergence and extension with constrained boundaries, cells invariably elongate parallel to the axis of convergence (Weliky et al., 1991; Brodland, 2006;). In contrast, when a tissue is passively strained along one axis, cells both elongate and separate through passive intercalation along the axis of loading (Keller and Trinkaus, 1987; Chen and Brodland, 2000; Aigouy et al., 2010). We have shown that in the fNP, cells both elongate and separate along the ML axis (Fig. 3d, e and h), perpendicular to the axis of convergence (Fig. 3g and Supplementary Table 3), with the signature of passive intercalation.

We wanted to understand how directionality could be imposed in the fNP, and what was the significance of the 5–7% average ML cell elongation associated with passive intercalation. We therefore set out a conceptual model of fNP dynamics, accounting for the mechanical properties of tissues surrounding the fNP, which we explored more formally in a cell-based simulation, that also gave us an estimate for the fNP of a key property of intercalating systems.

### A conceptual model of fNP dynamics

The anterior boundary of the fNP borders a large domain of non-neural ectoderm that expands in the plane during the contraction phase (Supplementary Fig. 6). The apposition of contracting neural and expanding non-neural domains allows the posterior displacement of their shared boundary. In contrast, lateral and posterior edges of the fNP border midbrain tissue, which is itself engaged in contractile neural morphogenesis and continuously connected by domains of neural tissue around the circumference of the embryos and pinned at the edge of the ectoderm (Fig. 2b).

To unify the morphogenetic signatures detected at multiple scales within the fNP, we propose the following model (Fig. 6). During the contraction phase, individual cells within the fNP actively contract in the plane, and approximately 50% undergo cell division (Supplementary Table 1), a process that facilitates radial cell rearrangements. Neither of these behaviours are planar-polarised and, in isolation, would effect isotropic and uniform planar tissue contraction, as observed in explants of amphibian fNP tissue (Jacobson and Gordon, 1976). We propose that *in vivo*, resistive tissue lateral to the fNP boundaries prevent its ML contraction. Consequently, planar cell intercalation emerges as a passive response to relieve tension generated by the ML component of contraction, thus converting isotropic contractile force into an anisotropic deformation that is uniform across the tissue. Finally, local radial compression by the PPM induces heterogeneity in the fNP deformation by transiently counteracting intrinsic cell behaviours in the medial region of contact.

**Figure 6.**
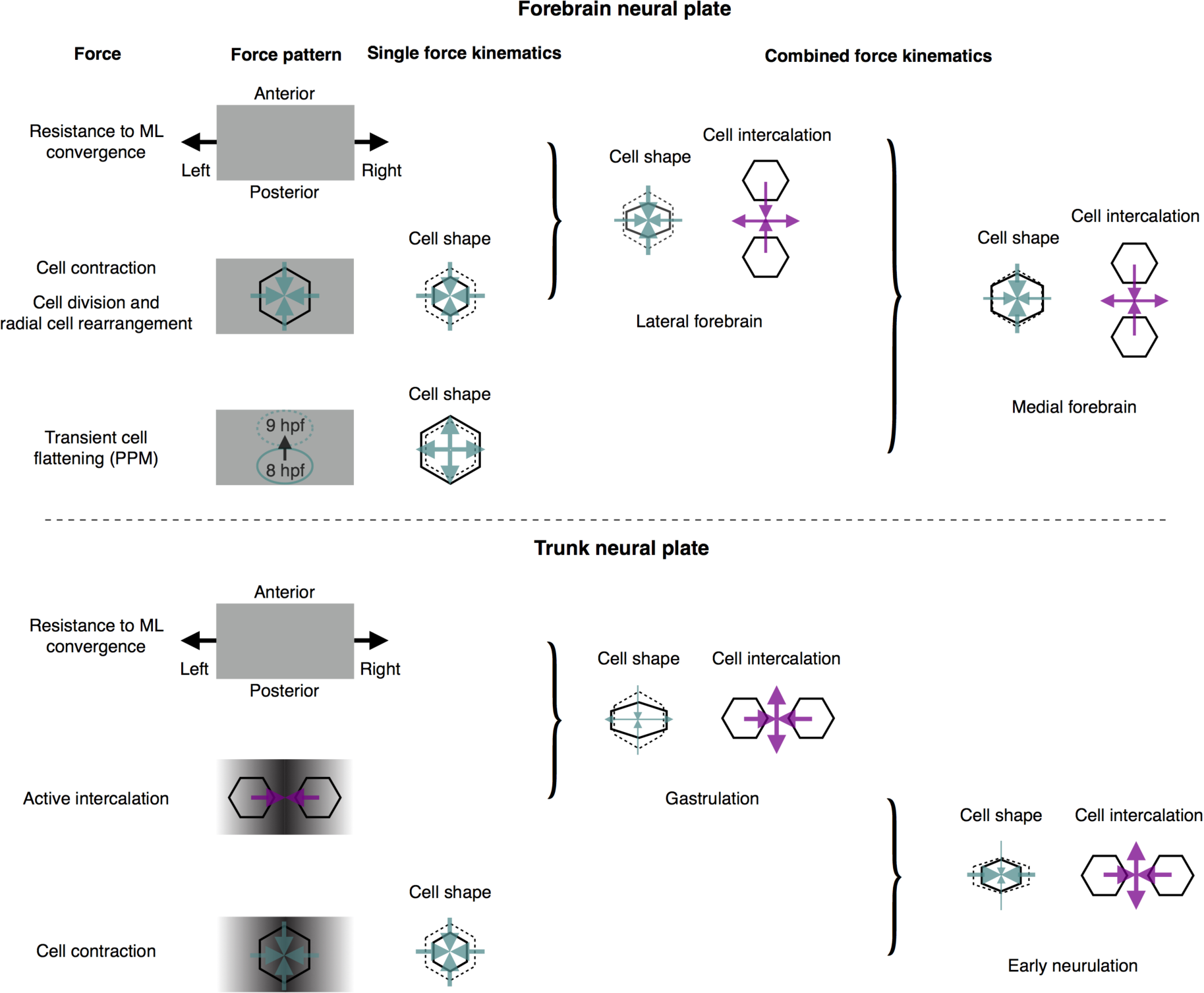
Comparing models of fNP and pNP cell behaviour and force combinations. A partially conserved set of cell behaviours and mechanical constraints result in comparable cellular deformations but distinct intercalation patterns and tissue deformations. Cells in the forebrain and trunk neural plate undergo planar contraction. Planar-polarised active intercalation in the trunk additively produces strong mediolateral tissue convergence while the forebrain fails to converge. Forebrain cells lacking planar-polarised behaviour instead passively separate along the mechanically-constrained ML axis.

To test the physical plausibility of this hypothesis, we developed a two-dimensional cell-based mechanical simulation of the fNP (see Methods) specifically designed to study the interplay between fundamental classes of active and passive cell behaviour with tissue boundary conditions. We first considered a uniform autonomous decrease in cell area in a tissue constrained by anisotropic boundary conditions representing the stiffness of the neighbouring tissues. This verified that isotropic cell contraction under anisotropic mechanical constraints leads to passive cell intercalation, wherein cells converge in the direction of lower stiffness (Supplementary Fig. 7).

We next considered whether observed cell shape and intercalation patterns in medial and lateral fNP could be a direct consequence of the pattern in cell area change (see Methods). Experimental time series of tissue area, for lateral and medial regions, were therefore used as model inputs (Fig 5, a and b, and Supplementary Movie 3); these implicitly capture intrinsic cell behaviours effecting area change, and medial compression by the PPM. Lateral tissue borders were also constrained to maintain the measured approximately stationary tissue width (Fig. 2g). We further assume that besides the intrinsic control of cell area, cellular properties are uniform across the tissue and constant throughout the simulation. No active intercalation or intrinsic cell asymmetry was programmed into the model. There are primarily two parameters left in the model to adjust the behaviour of the tissue, a cell intercalation relaxation time-scale, and a yield strain, the cell deformation at which plastic rearrangement begins. We have adjusted these parameters to fit the experimental data of the rates of cell shape change and of cell intercalation in the medial and lateral regions. We obtained close agreement between cumulative stretch ratios in the model and experimental data, within both lateral and medial regions (Fig. 5, e and f). Intercalation behaviour emerges here as a passive response (Supplementary Fig. 7). The fact that we reproduce the orientation and relative rates of cell shape deformation and cell intercalation within and between fNP regions from uniform material properties indicates that locally patterned or modulated mechanical responses are not necessary to produce the differential deformations in medial and lateral fNP regions. The values obtained for the model parameters provide estimates for the tissue rheological properties, with a yield strain in the range of 5% to 10%, and a time-scale of relaxation by intercalation of the order of 5 to 10 minutes. To test if these values are realistic, we plotted the relationship between cell elongation ratio and local intercalation rates for three embryos (Supplementary Fig. 8). Although the data is scattered, the trend is consistent with the model prediction, with yield strains in the range of the model and values for the relaxation time scale of approximately 30 minutes, which is larger than the model estimate but nevertheless of the right order of magnitude.

## Discussion

We provide the first quantitative description of cell and tissue dynamics during the earliest phase of forebrain neurulation in the zebrafish embryo. A number of unexpected and novel differences in fNP cellular behaviour relative to the spinal cord and hindbrain neural plate were identified. Our findings raise and support the novel concept that complex tissue movements of early forebrain morphogenesis do not require intrinsic polarised cell behaviours, but instead emerge from the structured mechanical landscape resulting from multiple interconnected and interacting tissues.

Early fNP neurulation is distinct from coincident pNP neurulation in three important respects. First, fNP cells intercalate within the plane to widen and shorten the tissue (Fig. 3, d and e, and Supplementary Fig. 3, b and c), whereas pNP cells do so to narrow and lengthen the tissue (Burnside and Jacobson, 1968; Jacobson and Gordon, 1976; Schoenwolf and Alvarez, 1989; Keller et al., 1992; Kimmel et al., 1994; Elul et al., 1997; Concha and Adams, 1998; Hirose et al., 2004; Hong and Brewster, 2006; Ybot-Gonzalez et al., 2007; Blanchard et al., 2009;). Planar intercalation has not previously been detected in the fNP. Our sensitive intercalation strain rate measure reveals this morphogenetic process that is essential for comprehension of cell movement patterns, and contributes to ~10% (lateral fNP) or ~20% (medial fNP) mediolateral tissue extension during the contraction phase in the zebrafish. Second, forebrain cells diverge radially to generate an additional cell layer from the initial two, while pNP cells converge radially to consolidate a single layer (Warga and Kimmel, 1990; Wilson et al., 1995; Kane et al., 2005; Hong and Brewster, 2006). Radial cell separation in the forebrain (Fig. 4d–i) is achieved predominantly by the movement of daughter cells following cell divisions. Increased tissue thickness and cell layer number is a recognised property of the fNP (Clarke, 2009), but here we offer this first mechanistic insight into how this local property develops. Despite random cell division orientation in the deep fNP cell layer (Figure 4j–m), radial separation of daughter cells may be actively controlled, possibly by asymmetric inheritance of an interface or tether to an emerging basal extra-cellular matrix. A comparable process was observed during later neural keel stages of hindbrain neurulation, in which one daughter cell maintains a basal tether and the other is displaced in the apical direction, although in this case cytokinesis itself is biased to the apico-basal axis (Tawk et al., 2007). In the surface fNP cell layer, cells possess more discoidal shapes with short axes perpendicular to the plane (Supplementary Fig. 1 and 2), cell division orientation is biased to the plane (Concha and Adams, 1998) (Supplementary Fig. 5, b and d), and radial separation of daughters is less stereotyped than in the deep cell layer (compare Fig. 4d, g to Fig. 4e, h). These differences between surface and deep layer cells may reflect differential interactions with apical and basal extra-cellular matrices respectively. The third difference between fNP and pNP neurulation is that cell divisions are not planar polarised in the fNP (Figure 4, j and l, and Supplementary Fig. 5, a and c), but highly oriented within the plane of the pNP (Concha and Adams, 1998), indicating that the fNP is not intrinsically planar polarised. Direct investigation of cellular polarity through measurements of molecular asymmetries in planar cell polarity proteins have yet to be performed in fNP tissue but will be of great interest.

Our analysis of cell behaviour across the fNP tissue leads us to propose a new conceptual model of tissue mechanics (see Results and Fig. 6). We were able to confirm the physical consistency of the key features of our conceptual model at the tissue and cellular level with a mechanical simulation consisting of isotropic cell contraction and anisotropic bounding tissue properties. We found a passive intercalation signature associated with cell elongation perpendicular to the orientation of neighbour gain. The precise elongation ratio of a cell and its associated local rate of passive intercalation contribute to the important tissue rheological parameters of yield strain and time-scale of relaxation by intercalation. Importantly, using our mechanical model we predicted values for these two rheological parameters and confirmed similar values from real data. The time-scale of relaxation by intercalation is short relative to the developmental time-scale of this morphogenetic process, and the yield strain was only modest at an elongation ratio of 5% along the ML cell axis. Given the concurrent active process of planar cell constriction, planar cell cortical tension (cell stiffness) is likely to be relatively high, which is an important prerequisite for relaxation by passive cell rearrangement on short time-scales and with modest yield strain. In contrast, other tissues that show zero or very slow intercalation even when cells become strongly elongated, such as the *Drosophila* wing disc (Mao et al., 2013) may be capable of cell rearrangement but lack the required cell stiffness.

Our results make strong predictions about the rheological properties and cell behaviours of the fNP and its surrounding tissues. Further testing of our model might include performing incisions along borders of the fNP or insertion of physical probes such as functionalised oil droplets (Campàs et al., 2014).

A comprehensive quantitative dissection of coincident events in the zebrafish pNP (G. B. Blanchard, N. L. Schultz, A. J. Kabla and R. J. Adams, Unpublished) also reveals a complex set of mechanical interactions whereby multiple intrinsic cell behaviours are tempered by the mechanics of surrounding tissues. Notably, both forebrain and neurulating trunk cells exhibit a comparable planar contraction process, as they become more columnar. However, cell contraction fails to effect mediolateral tissue convergence in the forebrain neural plate because adhesion between contracting cells is unable to sustain this stress against stiffer boundaries and stress dissipates by cells sliding apart passively. In the trunk neural plate, cell contraction initiates within a tissue already actively converging by planar-polarised active intercalation and able to exert a deformation on the surrounding ectoderm. In this case, planar contraction to columnarise the cells augments tissue convergence (Figure 6).

Finally, why should the fNP adopt such a derived behaviour? The region analysed is predominantly presumptive eye-field, which will be subsequently involved in substantial plastic rearrangement between two morphogenetic events: the deep and anterior-ward subduction of the hypothalamus and the dorsal-medialward convergence of the telencephalon (England et al., 2006; Cavodeassi et al., 2013). Tissue thickening and the relative ease with which its cells can passively rearrange would greatly facilitate this later remodelling, whilst preserving local tissue integrity.

Our ability to evaluate tissue-mechanical hypotheses pertaining to strain rate maps will improve as we accumulate examples of conserved relationships between orientations and quantitative contributions of interdependent cellular behaviours and the underlying *in vivo* stress fields and tissue-mechanical properties across diverse tissues. Encouragingly, it appears that the range of cellular behaviour combinations, and hence potentially the number of underlying mechanical morphogenetic modes, is not only finite but small.

## Methods

### Embryos

Wildtype embryos were ubiquitously labelled by injecting mRNA encoding farnesylated eGFP (plasma membrane marker, 180–200 pg) and histone-mCherry (nuclear marker, 80–90 pg) at the one cell stage. Embryos were dechorionated and mounted for live imaging as described (England et al., 2006).

### Imaging

Time-lapse sequences were recorded with a Leica TCS SP scanning confocal system mounted on an inverted microscope with a long working distance 20x/0.50 water immersion objective. Confocal stacks of 80–150µm in steps of 1–2.8µm were acquired at 2 or 2.5 minute intervals. Image series were deconvolved using the maximum-likelihood estimation algorithm of Huygens Essential (Scientific Volume Imaging B. V).

### Tracking

Custom manual and automated 2D tracking software in IDL (Exelis Visual Information Solutions) was used to reconstruct cell positions and outlines as previously (England et al., 2006; Blanchard et al., 2009). Automated 3D nuclear tracking was performed using a 3D instantiation of the previous 2D adaptive watershed segmentation and cell-linking algorithm, applied here to image stack volumes of the inverse of the histone signal. Trajectories were built connecting the darkest pixel of the smoothed histone signal for each cell at each time point. Automated 3D cell shape tracking was performed on movies with both nuclear and membrane channels, which were merged by adding the inverse of the nuclear channel to the membrane channel, after first coordinating the greyscale ranges of each channel. Cell shape tracking was then performed as for 3D nuclear tracking, storing the coordinates of the 3D watersheds as the cell shape outlines. Best-fit ellipsoids to these outlines were calculated for each cell at each time point for 3D cell shape and orientation analysis. Radial cell depth from the curved embryo surface for cells identified using all the above methods was defined as described (England et al., 2006; Blanchard et al., 2009).

### Strain rate measurement and decomposition into contributing cellular processes

Two-dimensional tissue strain rate tensors, estimating the planar deformation of overlapping tissue domains (Blanchard et al., 2009) from cell centroid movements were calculated through the contraction phase. The eigenvectors and eigenvalues of the tissue strain rate tensors represent the principal orientation and magnitude of tissue deformation. The domain size used for the calculation of tissue strain rates was 2 coronae (two ranks of immediate neighbours, ~70µm) and the time window was either +/-4 minutes or +/-5 minutes. We corrected for embryo curvature by projecting cell trajectories and cell shapes onto a plane tangent to the local surface of the embryo (Blanchard et al., 2009). Tissue strain rates were decomposed into a planar cell shape change strain rate (estimated directly from the evolution of cell outlines), a planar cell intercalation strain rate (that conserves area) and a residual area change due to cell division and radial cell rearrangement processes (assumed to be isotropic).

### Cumulative stretch ratios

Strain rates measured within selected tissue regions were projected onto the AP and ML axes of the embryo, averaged for each frame, integrated in time and exponentiated to give cumulative stretch ratio time series (Blanchard et al., 2009). To combine data from individual embryos with unequal time sampling, cumulative stretch ratio time series were interpolated and then averaged across embryos to produce a single average line. The endpoints of the original trajectories from individual embryos are shown to indicate the full range in the data.

### Planar cell area, orientation and elongation

Automatically tracked planar cell outlines in lateral and medial fNP regions were pooled from three independent embryos. 95% confidence intervals on the median cell area and projected cell elongation, ln(ML projected cell length/AP projected cell length), were calculated by bootstrap sampling using the *boot* package of R (R Development Core Team, 2011). The bin areas of cell long axis orientation radial histograms are proportional to frequency.

### Radial cell depth distributions

Radial cell depth time series (Fig. 4a–c) represent automated three-dimensional nuclear centroid tracking data from a single embryo. Radial cell depth analysis in Fig. 4d–m uses manually-tracked cell centroids. Kernel density estimates were constructed with the R *density* function using a Gaussian kernel (R Development Core Team, 2011). Sum of Gaussian models with one, two or three Gaussians were fit to kernel density estimates using custom software in R.

### Cell division orientation

Cell division orientation was defined as the three-dimensional vector connecting the centroids of daughter cell pairs at cytokinesis. This vector was defined manually for populations of cells representing lateral surface, lateral deep, medial surface and medial deep fNP tissue in three independent embryos; data were pooled for analysis. Cell division orientation distributions for each population (doubled, since undirected) were compared against a uniform circular distribution (representing a null hypothesis of randomly oriented cell divisions) at a critical value of p=0.05 using Watson’s one-sample goodness of fit test for circular uniformity in the R *circular* package (Concha and Adams, 1998; R Development Core Team, 2011).

### Simulation of the fNP

We constructed a cell based computational simulation of the fNP (Jennings, 2013). Each cell is modeled as an elastic domain with a shear modulus *G* that defines the resistance to changes in the shape of the cell while retaining a constant volume, and a bulk modulus *K* defining resistance to changes in the volume of the cell. Both moduli are constant over the course of simulations and are the same for all cells. The reference shape of each cell is circular, with a preferred area defined as one of the parameters of the model.

Cells interact with each other through mechanical forces. These forces are divided into two categories: contact forces and viscous forces. Contact forces capture the elastic force acting across the membrane due to cell compression and deformation. These forces have contributions from the cell internal pressure, cell mean deformation and local distortion, controlled respectively by three distinct parameters, K, B, and a parameter C that controls jamming in the system. Contact forces act normal to the interface between each pair of contacting cells. Viscous forces are associated with relative cell displacements at their interface and account for the molecular dynamics of cell adhesion. The parameter *η* defines the ratio between the viscous force and the sliding velocity of two neighboring cells.

From a configuration at a given time, the only unknowns are the viscous forces. Considering the physical constraints of force balance, torque balance and the fact that the internal stress of a cell has to be consistent with the distribution of forces at its interface, we can determine the viscous forces and the corresponding relative cell displacements that have to take place in order to satisfy mechanical laws. This allows us to update the tissue geometry and iterate for as long as necessary to follow morphogenesis on large time scales. A systematic analysis of the model behaviour shows that the emerging material properties are those of a visco-plastic material largely following the Bingham-Norton model (Irgens, 2008). Relaxation occurs in this model by cell intercalation, and the rate of intercalation 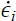 follows essentially the following relationship: 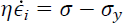 where *σ* is the shear stress in the tissue and *σ*_*y*_ the yield stress. For comparison with time lapse data, the relationship can be written in terms of strains 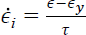. The yield strain at the cellular scale, *∈*_*y*_, is controlled by C. The relaxation time by intercalation, *τ*, scales like *η/G* (Jennings, 2013).

The initial preferred area of each cell is randomly selected from a distribution of cell areas measured from the initial state of the tissue *in vivo*. The active part of the dynamics is driven by specifying the preferred cell area as a function of time. We have introduced two separate programs for the time evolution of the average area of the cells within the medial (*Am(t)*) and lateral (*Al(t)*) sections of the tissue according to the actual experimental measurements of the cell areas in vivo (Fig. 5).

We first demonstrate how cell behaviour combined with anisotropic boundary conditions leads to intercalation in a simple model (Supplementary Fig. 7). In this model, all cells undergo a linear change to half their original area. The periodic boundaries behave as springs with stiffness *K*_*1*_ and *K*_*2*_ for the horizontal and vertical boundaries respectively. We vary the ratio of these stiffnesses between 1 and 20, and see that the larger the mismatch in stiffness, the more the system intercalates, converging along the soft direction. We set the timescale of the cell rearrangements (*τ*) to be fast compared to the timescale associated with the shrinkage, so the system is able to rearrange. The stiffness of the boundaries is made to be softer than the cells.

The model tissue is a periodic box wrapped in AP and ML. The time evolution of the length and width of the box are inputs supplied by the measured data. The system contains 512 cells and the initial aspect ratio of the box is 1.96. The cells in the middle third of the box are medial cells and all other cells are lateral. We started with *K*=20 and varied the timescale (*τ*) by varying the stiffness of the cells (*G*=0.5–2), the jamming parameter (C=10–40) and cell intercalation viscosity (*η*=20–40). We measured the distance between the *in vivo* data and the output data of the model to find the *τ* value that gave the best fit.

## Acknowledgments

A Wellcome Trust PhD Studentship awarded to S.Y., an MRC Project Grant awarded to R.J.A. and an EPSRC Project Grant awarded to R.J.A. and A.J.K. supported this research. We thank S. J. England for experimental advice and D. Bray, W. A. Harris, W. W. Lytton and J. Parker for comments on the manuscript.

## Author Contributions

R.J.A. and S.Y. designed the study. S.Y. performed experiments. S.Y. analysed data with input from G.B.B., A.J.K. and R.J.A. G.B.B. developed automated tracking software. The computational model was developed by S.Y., J.J., A.J.K. and R.J.A based on a platform created by J.J., A.J.K and R.J.A. S.Y. wrote the manuscript with R.J.A. All authors refined the manuscript.

